# Will Current Protected Areas Harbour Refugia for Threatened Arctic Vegetation Types until 2050?

**DOI:** 10.1101/2021.04.28.441764

**Authors:** Merin Reji Chacko, Ariane K.A. Goerens, Jacqueline Oehri, Elena Plekhanova, Gabriela Schaepman-Strub

**Affiliations:** Department of Evolutionary Biology and Environmental Studies, University of Zurich, Winterthurerstrasse 190, 8057 Zürich, Switzerland; Department of Environmental Systems Science, Institute of Terrestrial Ecosystems, ETH Zürich, Universitätstrasse 16, 8092 Zürich, Switzerland; Swiss Federal Research Institute WSL, Zürcherstrasse 111, 8903 Birmensdorf, Switzerland; Department of Biology, McGill University, Montréal, QC, Canada

**Keywords:** global change, arctic conservation, tundra vegetation, CAVM, vegetation shifts, climate change refugia

## Abstract

Arctic vegetation is crucial for fauna and the livelihoods of Northern peoples, and tightly linked to climate, permafrost soils, and water. Yet, a comprehensive understanding of climate change effects on Arctic vegetation is lacking. Protected areas cannot halt climate change, but could reduce future pressure from additional drivers, such as land use change and local industrial pollution. Therefore, it is crucial to understand the contribution of protected areas in safeguarding threatened Arctic vegetation types. We compare the 2003 baseline with existing 2050 predictions of circumpolar Arctic vegetation type distributions and demonstrate an overrepresentation of dominant and underrepresentation of declining vegetation types within protected areas. According to IUCN criteria, five of eight assessed vegetation types were classified as threatened by 2050. Potential climate change refugia, areas with the highest potential for safeguarding threatened vegetation types, were also identified. This study provides an essential first step to assessing vegetation type vulnerability based on predictions covering 46% of Arctic landscapes. The co-development of new protective measures by policymakers and indigenous peoples at a pan-Arctic scale requires more robust and spatially complete vegetation prediction, as increasing pressures from resource exploration and infrastructure development threaten the sustainable development of the rapidly thawing and greening Arctic.

## 1. Introduction

The Arctic experiences climate warming at twice the global mean (Collins et al., 2013; Walsh et al., 2011), leading to observed shifts in the distribution and composition of Arctic tundra vegetation (Serreze et al., 2000) that are projected to intensify in the future (Pearson et al., 2013a). These changes may threaten not only endemic plant species but entire vegetation types and the associated ecosystem functions. Vegetation types provide habitats for sessile and migratory animal species (Wheeler et al., 2018), support livelihoods of Northern peoples (CAFF, 2013) and take part in various feedbacks involving climate (Juszak et al., 2014; Loranty et al., 2011; Myers-Smith et al., 2011; Swann et al., 2010), permafrost soils (Bonfils et al., 2012; Juszak et al., 2014; Myers-Smith et al., 2011), lakes, rivers, and the ocean through water, energy and carbon fluxes (Lunt et al., 2004; Woo & Young, 2006).

Under the intermediate and high carbon emission scenarios (RCP 4.5 and 8.5), a recent long-term modelling study predicted that the current tundra area in Siberia will be reduced to 5.7% by 2500 (Kruse & Herzschuh, 2022). The southern border to the Arctic tundra is the treeline, which has been both observed and projected to advance northwards, decreasing the total extent of the tundra while simultaneously displacing southern Arctic vegetation types (Bjorkman et al., 2020; Chapin & Starfield, 1997; Lloyd et al., 2002; Rundqvist et al., 2011). Due to decreasing snow cover and phenological changes, primary productivity and vascular plant biomass have been increasing, specifically that of tall shrubs (Myers-Smith et al., 2011). This shrubification of the tundra has been widespread, resulting in a reduction in the abundance of less productive lichen- and moss-dominated vegetation types (Bjorkman et al., 2020; Blok et al., 2010; Elmendorf et al., 2012; Loranty & Goetz, 2012; Myers-Smith et al., 2011; Sturm et al., 2001). Widespread greening of the Arctic due to climate warming has long been identified (Myers-Smith et al., 2020; Myneni et al., 1997). However, spectral browning and increased heterogeneity of Arctic greening at the circumpolar scale have also been observed in recent years (Bjorkman et al., 2020; Phoenix & Bjerke, 2016), further complicating our understanding of changes in the highly heterogenous vegetation of the Arctic tundra.

Arctic vegetation is characterised by small vascular plants, bryophytes and lichens forming distinct plant communities. The zonal vegetation can be classified into five physiognomic classes (barrens, graminoid-dominated tundras, prostrate-shrub-dominated tundras, erect-shrub-dominated tundras, and wetlands), further divided into fifteen vegetation types based on the dominant plant functional types (Walker et al., 2005). The plant functional types are separated by plant traits such as growth form (e.g. graminoids, shrubs), size (e.g. dwarf or low shrubs), taxonomical status (e.g. sedges, rushes, grasses), and stature of woody shrubs. The resulting units include vegetation types such as cryptogram-dominated barren, graminoid-dominated, prostrate dwarf-shrub, low-shrub tundra, and sedge-dominated wetlands. They served as the fundamental units of the Circumpolar Arctic Vegetation Map (2003 CAVM) (CAVM Team, 2003), a project within the Conservation of Arctic Flora and Fauna (CAFF) group of the Arctic Council (CFG, 2022). Arctic vegetation types, as classified by the CAVM approach, relate to their ecological function in the Arctic system, using a standardised approach across the entire Arctic tundra biome. Hence, the CAVM approach has been widely applied to study very diverse processes relying on plant functional types, including biogeochemical fluxes (e.g. Virkkala et al., 2022; Zona et al., 2022), land surface energy flux (e.g. Oehri et al., 2022; Yu et al., 2022), land surface modelling improvements (Sulman et al., 2021), Alaskan tundra wildfire activity (Masrur et al., 2022) and studies investigating impacts of increasing shrub abundance on migratory songbirds (e.g. Boelman et al., 2015). However, a modelling study forecasted that 48–84% of Arctic vegetation types will have shifted by 2050 due to the effects of climate change (Pearson et al., 2013a).

Novel pressures on Arctic vegetation beyond those posed by warming are arising in the Arctic due to increasing anthropogenic presence. Historically, human land use and modification have been relatively low or non-existent in the Arctic biome. The human footprint map classifies most of the Arctic as under low pressure, and the human modification index demonstrates that the tundra remains thus far one of the last true terrestrial wild places on Earth (Kennedy et al., 2019; Venter et al., 2016). However, human interest in Arctic commodities such as oil and gas is increasing due to rising global energy needs, and with it also the extent of disturbances that terrestrial ecosystems experience due to exploration and infrastructure development (CAFF, 2013; Kumpula et al., 2011). The effects of human disturbances and landscape processes such as climate and biota shifts on the distribution and composition of vegetation over time must be explicitly included in effective conservation efforts. As vegetation in the Arctic is more fragile and requires longer times to recover from perturbation in comparison to southern vegetation (Kumpula et al., 2011), it is both an ecological, political and economic imperative to have plans in place to mitigate and minimise these disturbances in order to provide a path towards the sustainable development of the Arctic.

As of 2016, 20.2% of the terrestrial Arctic area is protected to some degree (CAFF & PAME, 2017). Although conservation efforts in the Arctic are well-developed, their focus is generally at the species level rather than the scale of ecosystems (Chapin et al., 2015). In other biomes, the impact of climate change-induced biota shifts on conservation efforts has been recognised, and adaptive conservation strategies have been developed, though not widely implemented (Heller & Zavaleta, 2009). A recent systematic study identified areas with high potential for the persistence of multiple biodiversity elements under climate change in North America, and demonstrated that at the biome-scale, ∼80% of areas within the top quintile of future conservation importance lacked formal protection, though this study excluded the High Arctic due to lacking data (Stralberg et al., 2020). Additionally, conservation needs in the North are influenced by rapid change at global scales and can no longer be addressed solely by local actions (Chapin et al., 2015). Indeed, local studies of Arctic plant communities do not always mirror observational trends over more extended temporal periods, which tend to be heterogeneous and complex (Bjorkman et al., 2020; Myers-Smith et al., 2020). Conservation actions which operate at various scales of space, time and biological organisation may be required to effectively prevent the loss of potentially vulnerable ecosystems and their functions.

Refugia have become a focus of interest in conservation, as they have been demonstrated to enable the persistence of biodiversity over longer temporal scales and changing climates while retaining ecosystem and habitat functions (Morelli et al., 2016; Tzedakis et al., 2002). Vegetation refugia can be defined as areas where existing vegetation will remain within current suitable climate conditions (Thorne et al., 2020). In this study, we consider refugia as areas where threatened vegetation types already occur, where the climatic envelope is predicted to remain suitable, and where competing vegetation types might still be absent due to dispersal limitations. Refugia may serve as a means for giving plant communities the time needed to allow for local adaptation to new environmental states by providing habitats where the effects of climate change are least felt in the short term (Morelli et al., 2020). The identification and protection of refugia may facilitate the persistence of retreating vegetation types under projected anthropogenic climate change (Noss, 2001; Taberlet & Cheddadi, 2002).

Presently, protected areas in the Arctic have not been established with climate change-induced vegetation shifts in mind (CAFF, 2013). Consequently, there is no consensus on the current state of vegetation vulnerability in the Arctic. Additionally, as Arctic vegetation shifts, today’s protected areas may no longer protect the same vegetation to the same extent in the future. Vegetation changes lead to trophic cascades, which alter the fundamental structures and functions of ecosystems (Wookey et al., 2009). Vegetation distribution projections for the year 2050 (Pearson et al., 2013b) provide the opportunity to locate vegetation refugia in the Arctic. Therefore, the conservation of potential vegetation refugia could, in effect, protect species at higher trophic levels. However, a comprehensive overview of vegetation refugia and their protected status in the Arctic has been unavailable to date.

This study aims to establish an overview of vegetation type abundance in Arctic protected areas and assess their vulnerability. For this purpose, we utilised the 2003 CAVM as the baseline abundance of Arctic vegetation types within and outside of current protected areas. We then determined the distribution of Arctic vegetation types within protected areas using previously existing 2050 vegetation scenarios (Pearson et al., 2013b). Furthermore, we assessed the risk of collapse of the vegetation types following the IUCN Red List of Ecosystems criteria for the baseline and the future (Bland et al., 2015). Lastly, to inform conservation efforts, we located potential refugia for the vegetation types that had been identified as threatened.

## 2. Materials and Methods

### 2.1. Baseline (2003) status of vegetation type abundance in protected areas

We defined the network of protected areas in the Arctic according to their extent in the Map of Arctic Protected Areas (MAPA) (CAFF & PAME, 2017). The protected areas were treated as one network regardless of the six management categories within the map, as our focus was on the collective pan-Arctic protection status. We utilised the 2003 CAVM to determine the baseline distribution of Arctic vegetation types within protected areas, which included fifteen distinct vegetation types (Appendix A) as well as the non-vegetative glacier class (CAVM Team, 2003; Walker et al., 2005). We intersected the 2003 CAVM with MAPA and computed a zonal histogram. This resulted in the pixel count of the protected area for each vegetation type in order to obtain the total and absolute areas of the vegetation types present within protected areas.

### 2.2. Future (2050) status of vegetation type abundance in protected areas

The bases for all analyses concerning future vegetation type distribution were the 36 Maps of Future Arctic Vegetation Distribution (MFAVD), which predicted climate change-induced vegetation shifts for 2050 (Pearson et al., 2013a). Two machine learning methods for ecological niche modelling, three climate models, two emissions scenarios, and three tree dispersal scenarios were applied to create 36 predictions (Pearson et al., 2013a). The MFAVDs are the result of all possible combinations of the parameters named above. Areas covered by glaciers, barren lands and wetlands were excluded from the predictions, but tree cover was incorporated to illustrate the northward treeline shift of the taiga. We intersected all 36 MFAVDs with MAPA to obtain the complete range of possible outcomes. We selected an MFAVD that we deemed to be most realistic (hereafter ‘realistic model’): random forest machine learning because it was the one with slightly higher accuracy (Pearson et al., 2013a); the intermediate of the three global climate models (i.e. CSIRO); the higher of the two emissions scenarios, A2a (IPCC, 2000); and the intermediate tree dispersal of 20 km, as it represents the highest rate at which trees have been observed to disperse northward in Alaska and Canada (Lescop-Sinclair & Payette, 1995; Lloyd et al., 2002).

We analysed the eight vegetation types available in the MFAVDs out of the fifteen originally present in the 2003 CAVM, as barren (four classes) and wetlands (three classes) had been excluded from the predictions (Pearson et al., 2013a). To assess the future distribution and protection status of Arctic vegetation, we converted MAPA from a polygon to a 4.5-km resolution raster to match the spatial resolution of the MFAVDs and attributed the protection status to every pixel. Finally, we summed the pixels and obtained the number of pixels per total as well as per protected area for each vegetation type.

### 2.3. Vulnerability of Arctic vegetation types following IUCN criteria

We applied the “IUCN Red List of Ecosystems Categories and Criteria” to quantify the vulnerability of Arctic vegetation types (Bland et al., 2015). The IUCN provides guidelines to assess the risk of ecosystem collapse, designed to be applicable to various ecosystems. Here, we defined ecosystems as the CAVM vegetation types, representing over 400 plant communities classified into broader functional groups (Walker et al., 2005). The IUCN assigns one of three graded categories to threatened ecosystems: “vulnerable”, “endangered”, and “critically endangered”. We did not differentiate here between the two graded categories for unthreatened ecosystems (“not threatened” and “least concern”) and instead simply summarised them as “not threatened”.

Five criteria are used to determine the IUCN risk category of the ecosystem: “reduction in geographic distribution”, “restricted in geographic distribution”, “environmental degradation”, “disruption of biotic processes or interactions”, and “quantitative analysis that estimates the probability of ecosystem collapse” (Bland et al., 2015). It is recommended that as many criteria as possible are assessed, as the ultimate classification of risk is determined as the highest of the five criteria (Bland et al., 2015). At the pan-Arctic scale, data were only available for “restricted distribution” and “decline of distribution”; therefore, this study could only assess these two criteria. Hence, our results represent the minimum risk status that each vegetation type can be assigned. Analyses of the remaining three criteria could potentially result in a higher risk category.

To determine the “restricted distribution” criterion classification, the number of 10×10 km grid cells occupied by the ecosystem must be calculated. An ecosystem is classified as “vulnerable”, “endangered”, or “critically endangered” if it occupies at most 50, 20 or 2 grid cells, respectively (Dudley, 2008). After the total extent of each vegetation type within the 2003 CAVM was calculated in square kilometres, the extents were divided by 100 to determine the number of 10×10 grid cells occupied by the vegetation type across the pan-Arctic extent. The “decline of distribution” criterion is determined by the predicted relative reduction in the distribution of an ecosystem over fifty years. An ecosystem is classified as “vulnerable”, “endangered”, or “critically endangered” if the reduction in its spatial extent is at least 30%, 50% or 80%, respectively (Dudley, 2008). In this analysis, the relative changes in area were calculated according to the changes between the 2003 CAVM (CAVM Team, 2003) and the 2050 MFAVDs (Pearson et al., 2013a); therefore, the classification is based on the predicted relative reduction over 47 rather than 50 years.

### 2.4. Identification of potential refugia for threatened vegetation types

For the threatened vegetation types, we identified areas which may serve as refugia, i.e. where a vegetation type is present in the baseline and is predicted to persist at least until 2050. We overlaid the realistic model with the 2003 CAVM and selected the areas where the pixels remained unchanged to create a map of areas with persistent vegetation types. We did not consider areas where these vegetation types had been predicted to shift to and only considered the areas where the vegetation remained the same, as a) the predictions of future vegetation distribution contain a measure of uncertainty, b) there is evidence of lag effects in species-scale responses to climate change (Stewart et al., 2016), and c) considerable variability in their ability to track new climatic niches exists (La Sorte & Jetz, 2012).

Spatial data were analysed using ArcMap 10.5.1 (ESRI, 2017) and RStudio 1.2.5033 (R Development Core Team, 2017; Team RStudio, 2020). Maps were converted into the Lambert Azimuthal Equal Area Polar Projection to preserve area, a crucial feature for this analysis.

## 3. Results

### 3.1. Baseline (2003) status of vegetation type abundance in protected areas

The terrestrial pan-Arctic region covers over 7.1 million km^2^, 20.2% of which is protected. Of the vegetated areas (4.7 million km^2^), 21% — or approximately 977,000 km^2^ — fall within protected areas. Additionally, a significant portion of the protected areas encompasses glaciated areas (777,000 km^2^) but could not be considered within the scope of this study due to a lack of predictions.

Our analysis demonstrates that the fifteen vegetation types have substantially different absolute and relative spatial abundances in the pan-Arctic tundra (Figure 1). The most abundant vegetation type (S1, erect dwarf-shrub tundra) covers over six times more area than the least abundant vegetation type (W1, sedge/grass, moss wetland), 626,000 km^2^, and 102,000 km^2^, respectively. The abundances of the vegetation types within protected areas generally mirror their abundance at the pan-Arctic scale. For example, 11.4% of Arctic vegetation is composed of the G3 vegetation type (nontussock sedge, dwarf-shrub, moss tundra), which accordingly covers 11.3% of protected areas. Notable exceptions do occur, such as the wetland types. In comparison to their total extents (W1: sedge/grass, moss wetland, 2.0%; W2: sedge, moss, dwarf-shrub wetland, 2.7%; W3: sedge, moss, low-shrub wetland, 3.7%), they are overrepresented within protected areas (W1, 3.0%; W2, 4.8%; W3, 6.7%). Contrastingly, though 7.5% of Arctic vegetation is composed of the vegetation type B2 (cryptogam barren complex, bedrock), it spans only 2.5% of the protected area. In terms of absolute extents, the most abundant types at pan-Arctic scales are overrepresented in protected areas, while the least abundant types are underrepresented.

**Figure 1.**
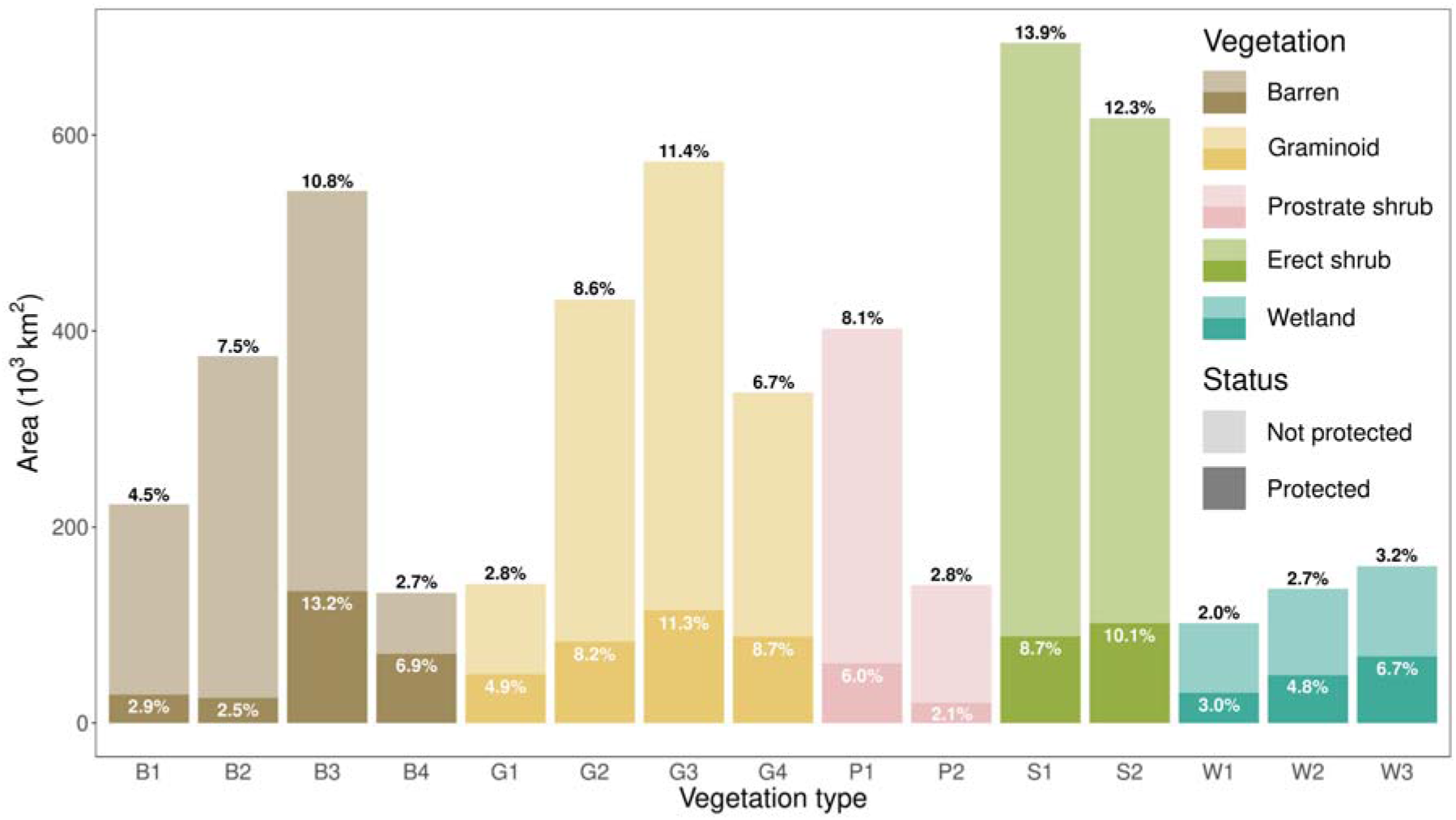
The 2003 baseline abundance and protection status of Arctic vegetation types. The Arctic tundra encompasses approximately 4.7 million km^2^. The height of the bars represents the absolute extent of each vegetation type within and outside of protected areas. The percentages above the bars represent the relative abundance of the vegetation type within the pan-Arctic tundra. The percentages within the darker coloured bars (protected area) represent the relative abundance of each vegetation type in the protected area in comparison to the total protected area. The vegetation types can be summarised into barren tundra (B1: cryptogam herb barren; B2: cryptogam barren complex (bedrock); B3: noncarbonate mountain complex and B4: carbonate mountain complex), graminoid tundra (G1: rush/grass forb, cryptogam tundra; G2: graminoid, prostrate dwarf-shrub, forb tundra; G3: nontussock sedge, dwarf-shrub, moss tundra; G4: tussock sedge, dwarf-shrub, moss tundra), prostrate-shrub tundra (P1: prostrate dwarf shrub, herb tundra; P2: prostrate/hemiprostrate dwarf-shrub tundra), erect-shrub tundra (S1: erect dwarf-shrub tundra; S2: low shrub tundra) and wetlands (W1: sedge/grass, moss wetland; W2: sedge, moss, dwarf-shrub wetland; W3: sedge, moss, low-shrub wetland).

### 3.2. Predicted vegetation type abundance in protected areas by 2050

Compared to the baseline abundances of vegetation types, the assessment of predicted abundances across protected areas by 2050 includes only eight vegetation types (see section 2.2). Predictions for the vegetation types within the barren (25.6% of non-glaciated terrestrial protected areas, 260,000 km^2^) and wetland (14.5% of non-glaciated terrestrial protected areas,148,000 km^2^) classes were unavailable. Additionally, as the nearly 777,000 km^2^ of glaciated protected areas were lacking in predictions, they were excluded from the study. The results differed considerably between the 36 MFAVDs: between 48,000 (0.1%) and 1,826,000 km^2^ (39.9%) of the total area covered by tundra vegetation in the 2003 CAVM is predicted to be replaced by taiga (Pearson et al., 2013a). Hence, considerable differences also resulted in the predictions of single vegetation types as a function of MFAVDs. We restrict the presentation of results to our selected realistic model and present the results of the other MFAVDs as an envelope around this model result. Of the eight vegetation types, five are predicted to decline in their total area according to the realistic model, accompanied by a mirrored decline in their abundance within the protected area network (Figure 2). In contrast, two southern vegetation types (S2: low shrub tundra and G3: nontussock sedge, dwarf-shrub moss tundra) are predicted to gain in area, both within and outside of protected areas. Vegetation type G4 (tussock sedge, dwarf shrub, moss tundra) showed a decrease in total area, yet its abundance within the protected area network increased.

**Figure 2.**
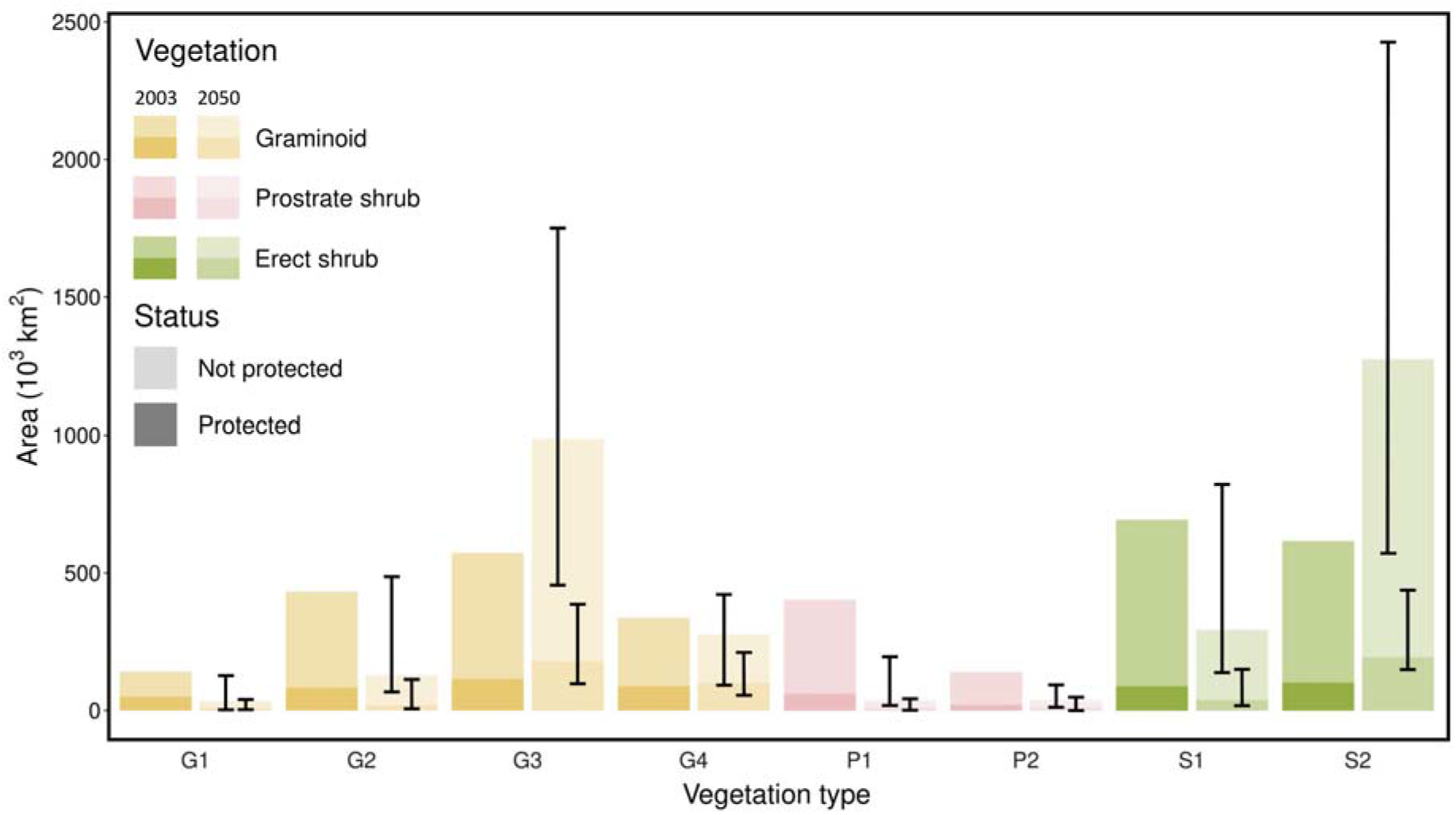
A comparison of the abundance and protection status of Arctic vegetation types at the baseline and in the future. The data presented refers to the realistic model scenario; error bars represent the envelope of the minimal and maximal abundance predicted by the different MFAVDs. The minimum and maximum values of the envelope are not always the result of the same MFAVDs. Depending on the vegetation type, different models predict the highest or lowest abundance. The vegetation types can be summarised into graminoid tundra (G1: rush/grass forb, cryptogam tundra; G2: graminoid, prostrate dwarf-shrub, forb tundra; G3: nontussock sedge, dwarf-shrub, moss tundra; G4: tussock sedge, dwarf-shrub, moss tundra), prostrate-shrub tundra (P1: prostrate dwarf shrub, herb tundra; P2: prostrate/hemiprostrate dwarf-shrub tundra), and erect-shrub tundra (S1: erect dwarf-shrub tundra; S2: low shrub tundra). Barren and wetland vegetation types were not assessed as predictions for 2050 were unavailable.

### 3.3. Vulnerability of Arctic vegetation types following IUCN criteria

To determine the risk of collapse for the eight vegetation types, we analysed two spatial criteria: “restriction of distribution” and “decline of distribution”; the final assigned category of risk was determined as the higher of these two. The “restriction of distribution” criterion diagnosed none of the eight vegetation types as threatened because each vegetation type has an extent of at least 5,000 km^2^. The “decline in distribution” criterion assigned different results (Figure 3). Depending on the MFAVD, the specific risk status assigned to each type varied. Under the realistic model, one vegetation type (P1: prostrate dwarf-shrub, herb tundra) was classified as critically endangered. Four vegetation types were classified as endangered (G1: rush/grass forb, cryptogam tundra; G2: graminoid, prostrate dwarf-shrub, forb tundra; P2: prostrate/hemiprostrate dwarf-shrub tundra; S1: erect dwarf-shrub tundra), and three as not threatened (G3: nontussock sedge, dwarf-shrub, moss tundra; G4: tussock sedge, dwarf-shrub, moss tundra; S2: low shrub tundra). The threatened vegetation types (i.e., those classified at least as endangered) are generally the more northern types with sparse, low-growing, non-vascular vegetation.

**Figure 3.**
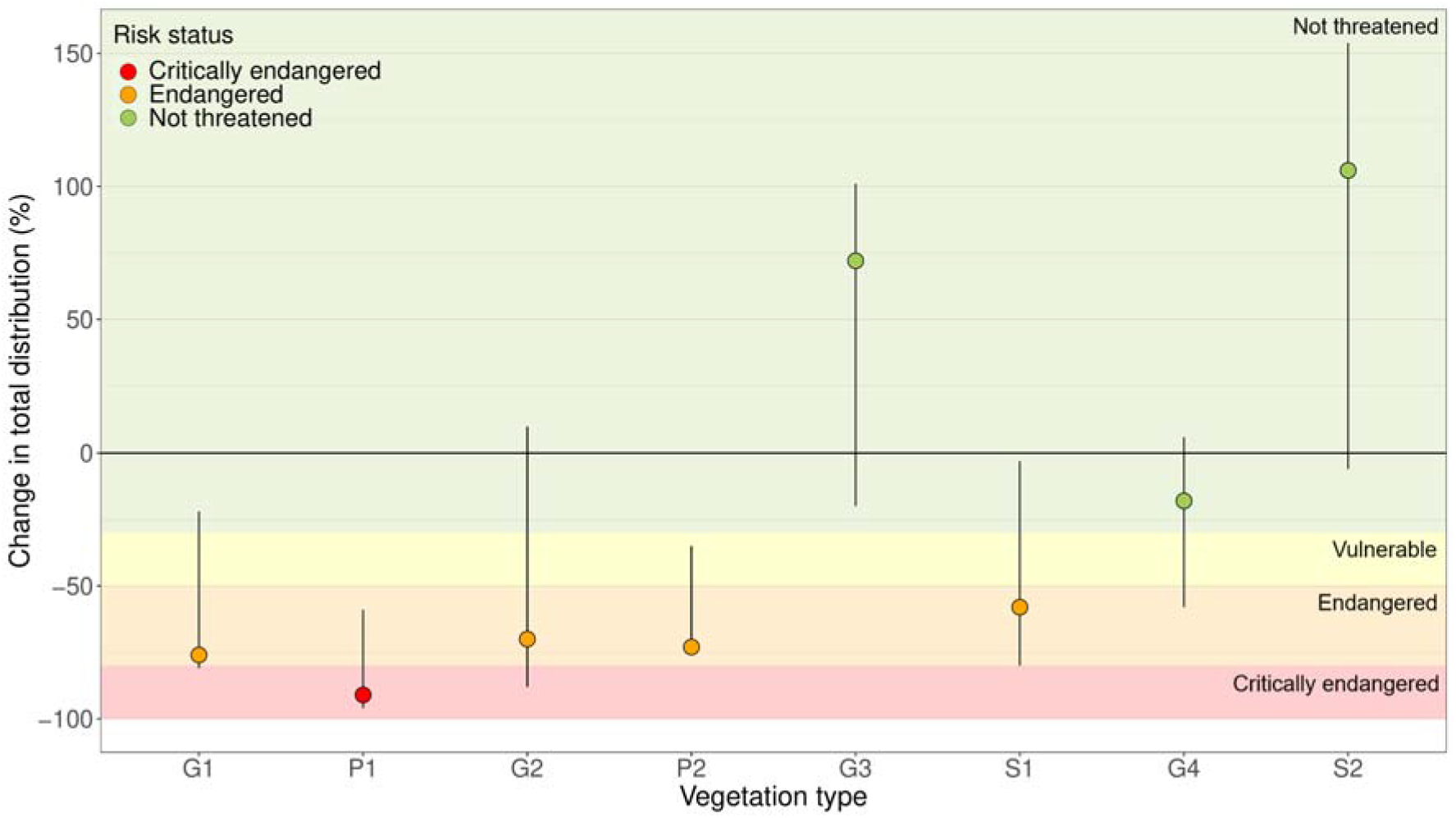
Risk status according to the ‘decline in distribution’ criterion of the IUCN Red List of Ecosystems Categories and Criteria under the realistic model. The vegetation types are ordered along the bioclimatic subzones they occur in, from low to high summer temperatures (left to right). The data presented refers to the realistic model scenario; error bars represent the envelope of the minimal and maximal abundance predicted by the different MFAVDs. The minimum and maximum values of the envelope are not always the result of the same MFAVDs. Depending on the vegetation type, different models predict the highest or lowest abundance. The vegetation types can be summarised into graminoid tundra (G1: rush/grass forb, cryptogam tundra; G2: graminoid, prostrate dwarf-shrub, forb tundra; G3: nontussock sedge, dwarf-shrub, moss tundra; G4: tussock sedge, dwarf-shrub, moss tundra), prostrate-shrub tundra (P1: prostrate dwarf shrub, herb tundra; P2: prostrate/hemiprostrate dwarf-shrub tundra), and erect-shrub tundra (S1: erect dwarf-shrub tundra; S2: low shrub tundra). Barren and wetland vegetation types were not assessed as no predictions for 2050 were available.

### 3.4. Potential refugia for threatened vegetation types

For the five vegetation types (G1, G2, P1, P2, S1) classified as threatened under the realistic model, we determined refugia, regions where these vegetation types are predicted to persist until 2050 (Figure 4, Appendix C). The total area of refugia ranges from < 9,000 km^2^ (P2) to > 100,000 km^2^ (S1), accounting for 2.1% and 8.7% of currently protected areas. The refugia are scattered over Canada, Greenland, Norway, Russia, and the USA (Figures 4a-d).

**Figure 4.**
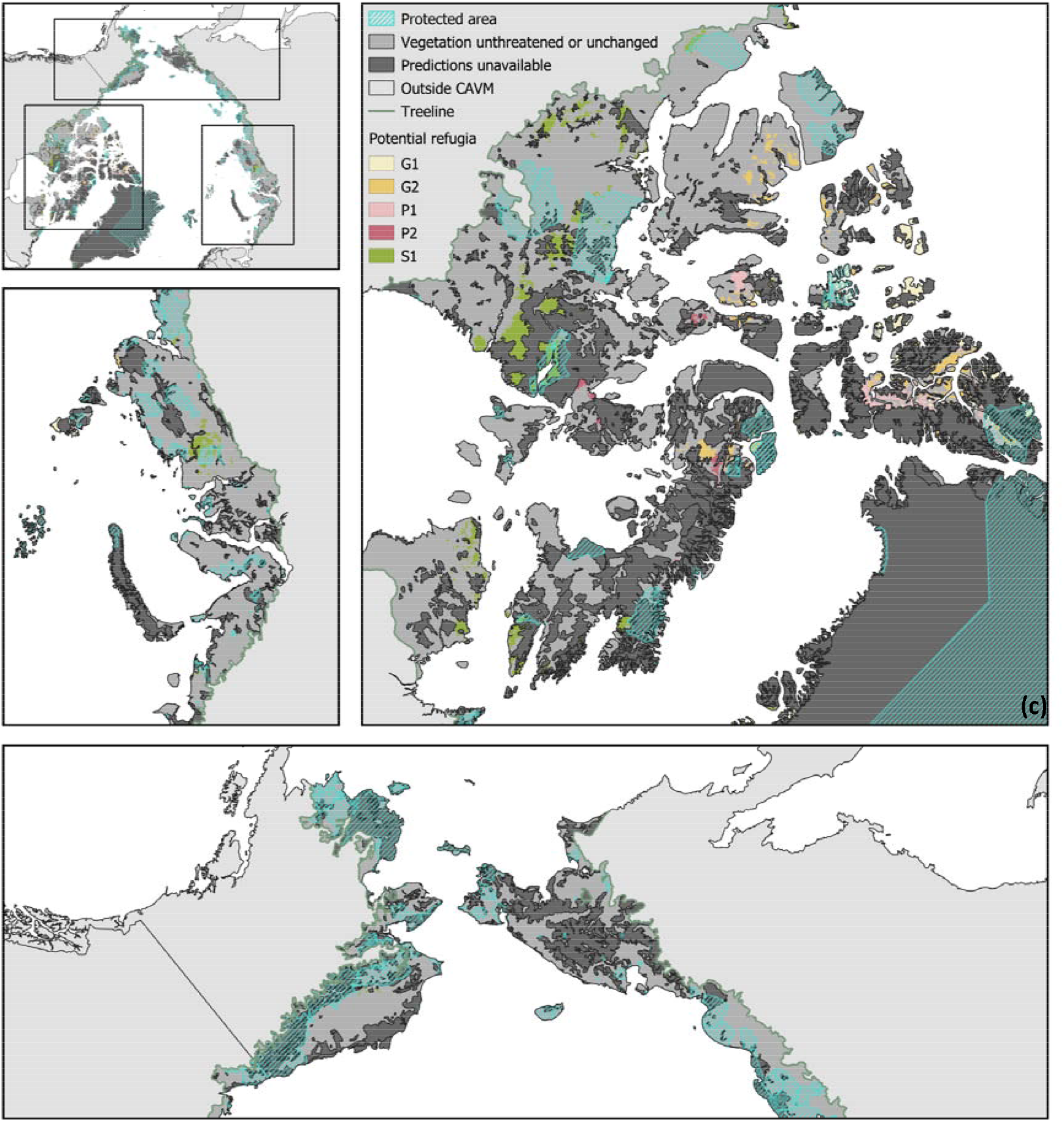
Refugia maps of threatened vegetation types in the Arctic (a), and magnified subsets centred on the Yamal peninsula, Russia (b); the Canadian Archipelago (c); and Alaska, USA, and North-eastern Siberia, Russia (d). The vulnerable vegetation types presented here are G1: rush/grass forb, cryptogam tundra; G2: graminoid, prostrate dwarf-shrub, forb tundra; P1: prostrate dwarf shrub, herb tundra; P2: prostrate/hemiprostrate dwarf-shrub tundra and S1: erect dwarf-shrub tundra.

## 4. Discussion

We evaluated the capability of the current Arctic network of protected areas for the conservation of Arctic vegetation types. Our results demonstrate that all Arctic vegetation types are found within the protected areas, and their relative abundance significantly varied but generally followed the same pattern as seen at the pan-Arctic scale. Noticeably, wetlands tend to be protected to an above-average extent; this is likely due to conservation efforts focusing on Wetlands of International Importance (Ramsar sites) related to their protection as breeding areas for migrating birds (CAFF & PAME, 2017).

The impacts of climate change may demand the adaptation of protected areas (Hannah et al., 2002; Heller & Zavaleta, 2009), as they may no longer harbour the same vegetation types. Thus, we analysed the protection and threat status of the vegetation type abundances predicted for the year 2050. The two most productive vegetation types (G3 and S2) occupying the lower southern bioclimatic zones were demonstrably increasing in extent. The five vegetation types which were predicted to decline drastically currently occupy the three more northern bioclimatic zones (Pearson et al., 2013b) and are characterised by sparse and low-growing vegetation (Raynolds et al., 2019). Our results demonstrate that these northern vegetation types also decrease in their representation within protected areas. We attribute this to two reasons. First, protected areas are generally situated in more southern regions rather than the northernmost edges of the Arctic, where these northern vegetation types can find sanctuary. This interpretation is validated by the fact that the refugia we located are clustered around the northernmost edges of the terrestrial Arctic. In our realistic model, we observe them being especially abundant in the Canadian Archipelago, which contains most of the terrestrial High Arctic. The climate will stay unsuitable for southern vegetation types in these regions at least until 2050. Secondly, Arctic protected areas were not established with the conservation of vegetation types in mind (CAFF, 2013); therefore, it stands to reason that vegetation types of conservation importance have not been recognised and hence prioritised.

### 4.1. Implications for flora, fauna and indigenous people if vegetation types are left unprotected

Vegetation shifts have been observed to have outcomes for species and communities across trophic levels (Myers-Smith et al., 2011; Wheeler et al., 2018). Threatened vegetation types may lead to the increased vulnerability of endemic flora and fauna, especially those that occur only in the High Arctic, which are most at risk. For example, there is evidence that vegetation exerts bottom-up control on caribou populations, as increasing shrubification decreases pasture quality (Fauchald et al., 2017). This decline is exacerbated by the loss of lichen-rich vegetation types, which negatively affect caribou populations due to a loss of winter foraging options (Cornelissen et al., 2001; Joly et al., 2009) and energy through increased methane emission if lichens are reduced in their diet (Hansen et al., 2018). Indigenous knowledge also shows that caribou populations have already adopted more northern migration patterns (Ksenofontov et al., 2019). This can, in turn, have negative consequences for indigenous communities that depend on caribou for food and as economic resources (Joly et al., 2009; Nuttall et al., 2005). Without focused conservation efforts on maintaining these vegetation types, the species depending on them will also become threatened with extinction.

### 4.2. Implications of land use change on threatened vegetation types if left unprotected

Climate warming and technological advances have opened up the Arctic as a new frontier in economic development. Increasing interest in commodities such as oil, gas, and mineral resources will naturally lead to increased infrastructure requirements in the Arctic, which may, in turn, intensify disturbances in Arctic ecosystems. Surface disturbances such as road networks and settlements have led to permafrost degradation through soil warming (Forbes et al., 2001). Off-road vehicles like tundra tractors can destroy endemic vegetation and leave the tracks visible for decades as vegetation struggles to recover (Forbes et al., 2001). A study of the Alaska North Slope demonstrated that 34% of the area was affected by oil development by 2010 (Raynolds et al., 2014). Further exploration has already been proposed, such as a 3-D seismic survey covering 63,000 km of trails within the Arctic National Wildlife Refugia in Alaska (SAExploration, 2018). A recent study demonstrated that this could result in mid-to-high level impacts on 122 km^2^ of the area in question, leading to increased thermokarst formation and erosion, and negatively impacting moist vegetation types (Raynolds et al., 2020). Additionally, vehicle tires can significantly increase the dispersal distance of southern invaders, such as *Salix lanata*, which has been found to occur in many roadside areas, replacing lichen and moss cover (Raynolds et al., 2014). Local warming due to the settlements also generates pockets of suitable habitats in latitudes where southern vegetation could not otherwise persist (Forbes et al., 2001).

Permafrost thaw is critically threatening Arctic infrastructures (Hjort et al., 2018), and their collapse can lead to dramatic consequences for terrestrial ecosystems and indigenous livelihoods. As the MFAVDs did not account for human land use modification, this study severely understates their effects on vegetation shifts. At the pan-Arctic scale, a combination of economic, geopolitical, climatic, infrastructure and ecological factors lead to uncertainty in the future spatial distribution of development pressures and their degree of impact (Scibilia et al., 2015). Therefore, it is essential to predefine conservation areas for the currently unprotected and threatened vegetation types before widespread development begins in the increasingly accessible Arctic. Complementing protected areas, biodiversity conservation schemes developed for other regions of the world, such as around mining sites, need to be adapted to Arctic conditions to prevent loss of vegetation where industrial development outside of protected areas is allowed.

### 4.3. Implications of using climate change refugia for land conservation and management

Globally protected areas are 10.6% richer in species diversity than non-protected areas (Gray et al., 2016). Nevertheless, conserving biodiversity using protected areas poses to be difficult under climate change. A study assessing the protected areas of Canada found that climate change will lead to over 40% of the protected areas experiencing a change in biome type, with the total extent of the Canadian tundra standing to decline by 38–79% (Lemieux & Scott, 2005). In order to most efficiently use limited resources and protect threatened vegetation, efforts in in situ management must be focused on areas where the vegetation is likely to persist under global change, such as climate change refugia for vegetation. There is scepticism over the use of refugia as areas where vegetation may retreat until conditions in the surrounding environment become more favourable. Vegetation shifts due to climate warming are projected to continue and intensify; therefore, the time periods required until favourable conditions return may be longer than the existence of these refugia. However, they may serve a powerful purpose by buying time for the climate adaptation of vulnerable species and ecological communities (Morelli et al., 2020). Adjusting Arctic protection efforts in mitigating the consequences of global change with a focus on relatively transient climate change refugia may aid in a long-term transformation of these threatened vegetation types into novel community assemblages (i.e. ecological replacements) that have adapted to these environmental changes while performing the same ecological and habitat functions (Morelli et al., 2020). This process has been documented in the late Quaternary (Jackson & Overpeck, 2000).

### 4.4. Limitations and urgent need for comprehensive predictions of Arctic vegetation type distribution

The aim of this study is to construct a first assessment of the protected status of Arctic vegetation types in light of climate change based on existing vegetation predictions. These predictions (Pearson et al., 2013b) were based on the 2003 CAVM, a vector map with polygons of 14 km minimum diameter (Walker et al., 2005); thus, for the sake of consistency, it was utilised by this study instead of the more recent raster version (2019 CAVM) that is resolved at 1 km (Raynolds et al., 2019).Nevertheless, these disparities did not affect the results regarding the baseline risk status of the vegetation types; the general pattern of abundant vegetation types being better represented within protected areas holds, regardless of the map used (Appendix B).

Increasing spatial resolution in future predictions could reveal pockets of vegetation occurring outside of the main macroclimatic niche of a vegetation type. Such pockets could accelerate the spread of vegetation types into surrounding areas should the climate become more favourable. Potentially, southern vegetation types could then disperse northward at faster rates than modelled (Pearson et al., 2013a), increasing the displacement of northern vegetation types and potentially our risk classifications. This displacement may be balanced by potential northern micropockets, which would increase the total extent of refugia. Nevertheless, these disparities did not affect the results regarding the present risk status of the vegetation types. Though the “decline in distribution” criterion could not be tested as MFAVDs do not exist for the 2019 CAVM, none of the vegetation types of the 2019 CAVM showed a restricted distribution and were classified as unthreatened under the IUCN “restriction of distribution” criterion.

Due to limited data availability in the Arctic, we could only analyse two of the five IUCN criteria. Therefore, the assigned risk category only represents the minimal risk and could potentially be higher, both in the present and in the future. Though it is recommended that as many criteria as possible are assessed, the IUCN Red List of Ecosystems was created with the purpose of flexibility in the use of data containing variations in quality and coverage (Bland et al., 2015). Therefore, as data availability increases, this risk assessment still provides valuable information as a starting point for future assessments.

The MFAVDs only rendered predictions for 46% of the pan-Arctic area. Specifically, they excluded glaciers (including nunataks), barren lands, and wetlands (Pearson et al., 2013a). Subsequently, we were unable to assess the future distribution of wetland and barren vegetation types, classify their risk of collapse, and locate potential refugia. Therefore, our results do not consider the potential of the assessed vegetation types to disperse into areas currently covered by these unassessed vegetation types and vice versa.

Glaciers currently cover nearly one-third of the Arctic (Walker et al., 2005), but extensive climate change-induced glacier melt is predicted (Alley et al., 2010; Hinzman et al., 2005). Vegetation succession following retreating glaciers in the Arctic follows the same patterns as elsewhere; however, these processes occur at larger temporal scales and may be less relevant for the next thirty years (Hodkinson et al., 2003; Raynolds & Walker, 2009). Vegetation growth response to warming is slower in the High Arctic than in the Low Arctic due to lower growing season temperatures, seasonal length and nutrient availability (Walker et al., 2006). In Svalbard, for instance, vascular plant establishment was highly limited for the first century after glacial retreat (Hodkinson et al., 2003).

Additionally, vegetation can take up to 500 years to reach equilibrium after glacial retraction in Greenland (Lunt et al., 2004). Therefore, the inclusion of glaciers would have likely had limited impacts on the results for threatened vegetation types shown here for 2050. Nevertheless, they would be crucial for imminent studies which investigate the future of the terrestrial Arctic at longer timescales.

One-quarter of the terrestrial Arctic consists of barren lands (Walker et al., 2005), which were excluded in this analysis. Therefore, the northern vegetation types may be less threatened than indicated by this study because they may expand into currently barren areas. Including barren lands in future studies would likely lead to predictions of northern vegetation types being slightly more abundant and increasing over centennial periods. However, due to the slow response in the development of soils and vegetation, the time scales at which this northern expansion into barren lands occurs may be longer than the 47-year span of our study, and this expansion may be further limited as barren lands lack in soil sufficient enough to support vegetation (Pearson et al., 2013a).

Wetlands constitute 7% of the Arctic vegetated area (Walker et al., 2005). Predictions for future wetland distribution are particularly difficult due to the complexity of the hydrological processes regulating them (Walvoord & Kurylyk, 2016). Permafrost thaw, increasing precipitation, ice melt, and evaporation due to warming influence the abundance and distribution of surface water, indirectly affecting wetlands (White et al., 2007; Woo & Young, 2006). In the southern areas with discontinuous permafrost, ponds are generally in decline as permafrost thaw increases drainage (Riordan et al., 2006; Smith, 2005; Yoshikawa & Hinzman, 2003). Conversely, the thawing of continuous permafrost in the High Arctic has increased abundance of thaw ponds (Smith, 2005). In the Canadian Arctic, this trend of thermokarst pond formation has been shown to occur at rapid paces (Farquharson et al., 2019). This pattern of wetland development means that generally, southern vegetation types would have more potential areas of expansion, whereas northern areas may become further displaced by expanding wetland. We hypothesise that the inclusion of wetlands in this study could consequently have a reinforcing effect on the observed patterns of threatened vegetation types. While wetlands are crucial because of the abundance of migratory birds dependent on them, they make up a relatively small portion of the vegetated pan-Arctic and are generally better protected than other vegetation types. However, this may change significantly in the future due to climate change. Our study highlights the need for further focus on these ill-understood vegetation types in longer-term predictions.

The future vegetation shifts (Pearson et al., 2013a) were predicted under various assumptions about climate change and tree dispersal rate scenarios, which, though necessary, carry over to this study. Uncertainties in future vegetation distributions are inherent to predictions based on climate models with many uncertain assumptions, where one model is no more or less valid than the other (IPCC, 2000). Our selection of the realistic future model with regard to climate and emission scenario is, therefore, subjective. There is a large spread of the vegetation type abundances across the models, making the results, especially for vegetation types G1, G2, G4 and S1, more uncertain (Figure 2 and Appendix D). However, the general trend of increase in southern vegetation types and decrease in northern vegetation types still holds within and outside of protected areas, as demonstrated for most results by the minimum and maximum scenario envelope.

While land surface schemes for earth system models and vegetation development models are being adapted to Arctic tundra conditions, parameterisations at the Arctic tundra vegetation type level, including all vegetation types, need to be implemented, and biotic and abiotic interactions integrated with holistic modelling approaches, at pan-Arctic scale. Furthermore, the inclusion of human land use scenarios (e.g. reindeer herding densities, potential mining sites, road development) will be necessary for future modelling efforts to allow for informed policies and decisions.

## 5. Conclusions

This study identifies baseline and future abundances of Arctic vegetation types within protected areas as well as potential climate change refugia for threatened vegetation types using the 2003 Circumpolar Arctic Vegetation Map and existing predictions of future Arctic vegetation distribution.

Though uncertainties exist within the maps provided here, the general trends seen in this study are valuable in guiding future conservation efforts. Further studies on the development of Arctic vegetation types which include the whole Arctic — especially wetlands, barrens and ice-covered areas — are urgently needed. They could provide complete and precise findings to improve and adapt conservation efforts to climate change-induced biota shifts. Moreover, we recommend independent validation of these hypothetical predictions to better incorporate climate change refugia into designated protected areas. This could potentially be achieved through recent validation methods which compare predictions of refugia to local measures of species richness, endemic species persistence, genetic diversity, plant functional traits and/or demographic variables (Barrows et al., 2020).

This study aims to shift the focus of pan-Arctic protection efforts towards higher scales of biological organisation, from species to communities and ecosystems, or in this case, vegetation types. Endemic tundra vegetation types face the threat of ecosystem collapse due to global change. The establishment of refugia for the vegetation types identified here could protect them as refugia have done over the history of life on earth (Alley et al., 2010; Jackson & Overpeck, 2000). The protection of vegetation refugia would additionally have positive effects on Arctic fauna and climate regulation through bottom-up biotic and abiotic interactions. However, the integration of vegetation types for climate change-adapted conservation in the Arctic requires urgent collaboration between policymakers and indigenous peoples as the area becomes increasingly under pressure from exploration and rapid infrastructure development. As experienced in the recent decade, Arctic change is rapid. Extreme events, including extreme winter precipitation and summer drought, are already affecting Arctic ecosystems through major disturbance events such as extensive flooding and fires, adding yet another dimension of abrupt change to mitigate in the future. Without a plan already in place for the protection of these critically important Arctic landscapes, we cannot enable the sustainability of economic and structural development in what is increasingly no longer one of the world’s last truly wild places.

## Supporting information

Appendices_Summary

## 6. Acknowledgements

This work was supported by the University Research Priority Programme Global Change and Biodiversity of the University of Zurich.

