## Appendices_Summary for "Will Current Protected Areas Harbour Refugia for Threatened Arctic Vegetation Types until 2050?"

**Appendix A:** The vegetation type classification of the Circumpolar Arctic Vegetation Map (CAVM)

The CAVM displays the types of vegetation that occur across the terrestrial Arctic, from the Arctic Ocean in the north to the boreal treeline in the south (CAVM Team, 2003). The goal of the CAVM team was to map zonal vegetation, i.e. vegetation that develops over time on mesic soils in balance with the local climate (Walker et al., 2005) in order to effectively summarise the general distribution of arctic vegetation (Raynolds et al., 2019). The vegetation is first classified into five broad physiognomic categories, namely, barrens (B, four classes) graminoid-dominated tundra (G, four classes), prostrate, dwarf-shrub-dominated tundra (P, two classes), erect dwarf-shrub-dominated tundra (S, two classes) and wetlands (W, three classes). Barren classes are predominantly barren soils or bedrock or covered by biological soil crusts characterised by a lack of much vascular plant cover (Raynolds et al., 2019; Walker et al., 2005). Graminoid tundra are areas dominated by graminoid plants (sedges, grasses and rushes). Prostrate-dwarf-shrub tundra areas are dominated by prostrate-dwarf shrubs. Erect-dwarf-shrub tundra areas are dominated by erect-dwarf shrubs (<40 cm tall) or low shrubs (40–200 cm tall) and mosses. Wetland complexes are predominantly composed of wet tundra, often with many lakes and ponds, and patterned ground associated with periglacial landforms (Raynolds et al., 2019; Walker et al., 2005).

**Appendix B:** The distribution of vegetation types within Arctic protected areas according to the new raster version of the CAVM.

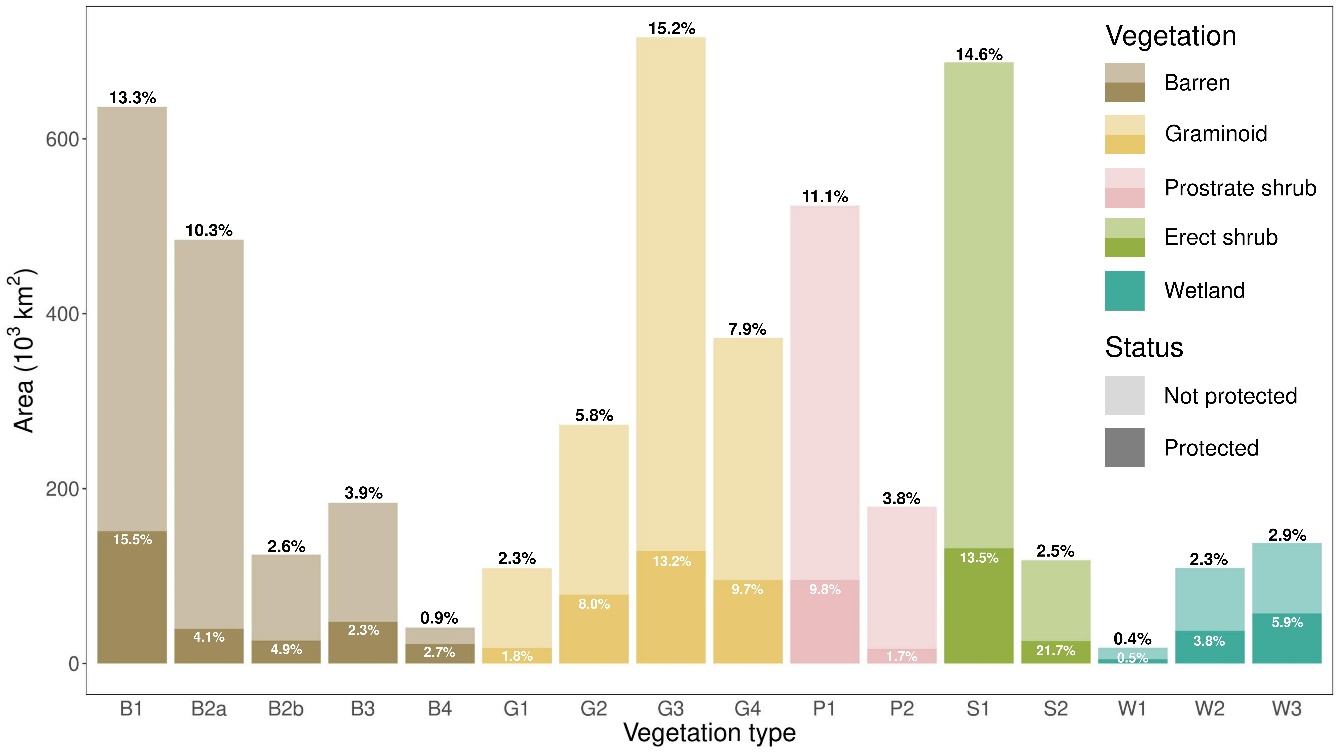

*Figure A.1.* **2019 CAVM abundance and protection status of Arctic vegetation types.** The height of the bars represents the absolute extent of each vegetation type within and outside of protected areas. The percentages above the bars represent the relative abundance of the vegetation type within the pan-Arctic tundra. The percentages within the darker coloured bars (protected area) represent the relative abundance of each vegetation type in the protected area in comparison to the total protected area. The vegetation types can be summarised into barren tundra (B1: cryptogam herb barren; B2a: cryptogam barren complex; B2b: cryptogam, barren, dwarf-shrub complex; B3: noncarbonate mountain complex and B4: carbonate mountain complex), graminoid tundra (G1: rush/grass forb, cryptogam tundra; G2: graminoid, prostrate dwarf-shrub, forb tundra; G3: nontussock sedge, dwarf-shrub, moss tundra; G4: tussock sedge, dwarf-shrub, moss tundra), prostrate-shrub tundra (P1: prostrate dwarf shrub, herb tundra; P2: prostrate/hemiprostrate dwarf-shrub tundra), erect-shrub tundra (S1: erect dwarf-shrub tundra; S2: low shrub tundra) and wetlands (W1: sedge/grass, moss wetland; W2: sedge, moss, dwarf-shrub wetland; W3: sedge, moss, low-shrub wetland).

Raynolds et al. (2019) produced a new raster version of the CAVM, with a higher resolution and better accuracy than the original CAVM. However, as the Maps of Future Arctic Vegetation Distribution (MFAVD) by Pearson et al. (2013) were created using the original CAVM, so did this study. Figure A.1. displays the current status of these vegetation types according to the new CAVM. Though the vegetation classes generally remained the same, B2 vegetation type was split into B2a and B2b. We evaluated differences in vegetation type abundance between the two CAVM versions for the baseline status to evaluate potential implications of using the 2003 CAVM. A Landsat accuracy assessment of the two maps shows that the 2003 CAVM had a 39% accuracy in comparison to 70% for the 2019 CAVM(Raynolds et al., 2019). The 2019 CAVM shows an increase in current P1 abundance both within and outside of protected areas in comparison to the 2003 CAVM (Appendix A). Therefore, the decrease in future abundance and threat may have been overestimated. Similarly, as the total extent of S2 decreased significantly from the 2003 CAVM to the 2019 CAVM, its future threat status may have been underestimated. This disparity between the two maps can be attributed to the lower resolution of the 2003 CAVM, which cannot capture the fine heterogeneity of Arctic vegetation type distribution (Raynolds et al., 2019).

**Appendix C:** Map of Potential 2050 Refugia for Vulnerable Arctic Vegetation Types

A map of potential refugia for the vulnerable arctic vegetation types discussed in this paper has been made available in the geospatial portable document format (Geospatial PDF), which preserves GIS functions such as layers.

**Appendix D:** Uncertainties in the usage of the ‘realistic model’ for the identification of refugia.

This paper focuses on the realistic model (random forest machine learning, CSIRO global climate model, the A2a emissions scenario, and the intermediate tree dispersal rate), which is one of 36 Maps of Future Arctic Vegetation Distribution (Pearson et al., 2013). However, as stated in the discussion, the spread of the data between vegetation type abundances and distributions across the models (figure 2). Therefore, to account for this uncertainty in the Map of Potential 2050 Refugia for Vulnerable Arctic Vegetation Types (Appendix C), a map which compares the vegetation classification in each pixel of refugia with the other 35 model vegetation predictions was made available in a portable document format (PDF).
