## Appendix_C for "Will Current Protected Areas Harbour Refugia for Threatened Arctic Vegetation Types until 2050?"

### Potential 2050 Refugia for Vulnerable Arctic Vegetation Types

#### Potential refugia

- Rush/grass forb, cryptogam tundra (G1)
- Graminoid, prostate dwarf-shrub, forb tundra (G2)
- Prostrate dwarf-shrub tundra (P1)
- Prostrate/hemiprostrate dwarf-shrub tundra (P2)
- Erect dwarf-shrub tundra (S1)
- Protected area
- Predictions unavailable
- Vegetation unchanged or unthreatened
- Treeline

WGS 1984 North Pole LAEA Bering Sea Projection

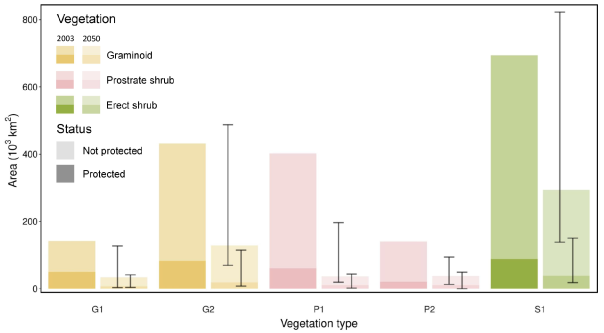


A comparison of the abundance and protection status of vulnerable Arctic vegetation types between 2003 and 2050.

**Citation**
M. Reji Chacko, A.K.A. Goerens, J. Oehri, E. Plekhanova and G. Schaepman-Strub. Will current protected areas harbour refugia for threatened Arctic vegetation types until 2050? *Manuscript submitted* .

**Project Director**
G. Schaepman-Strub

**Compilation and Cartography**
A. K.A. Goerens, M. Reji Chacko

University of Zurich[UZH]

URPP Global Change and Biodiversity

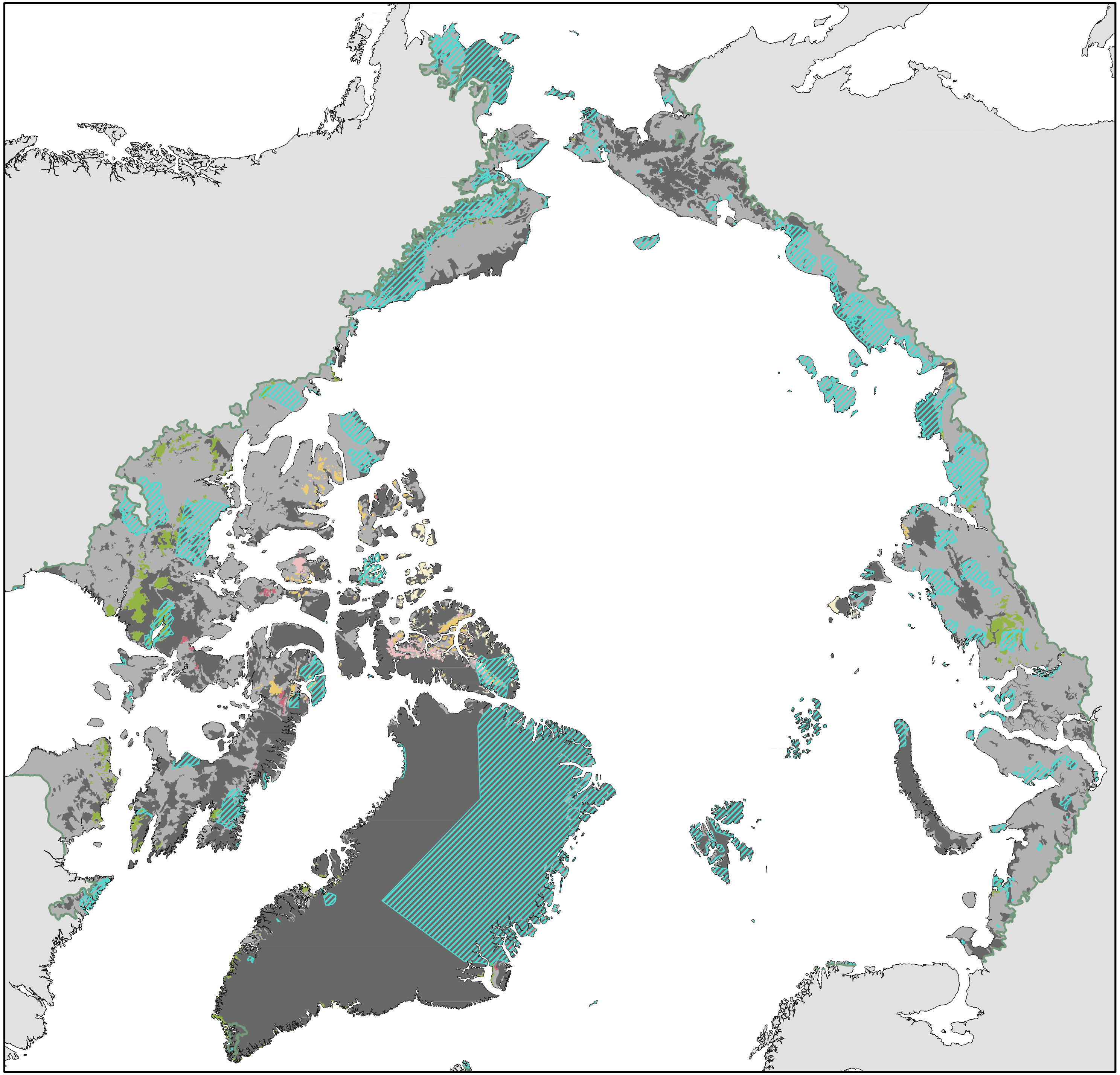
