## Appendix_D for "Will Current Protected Areas Harbour Refugia for Threatened Arctic Vegetation Types until 2050?"

### Uncertainties in Refugia Identification Under the Realistic Model Scenario

Frequency of other vegetation models which predict the same persistent vegetation class

- 1 - 9
- 10 -19
- 20 - 27
- 28 - 35

- Non-Arctic areas
- Arctic areas
- Treeline

WGS 1984 North Pole LAEA Bering Sea Projection

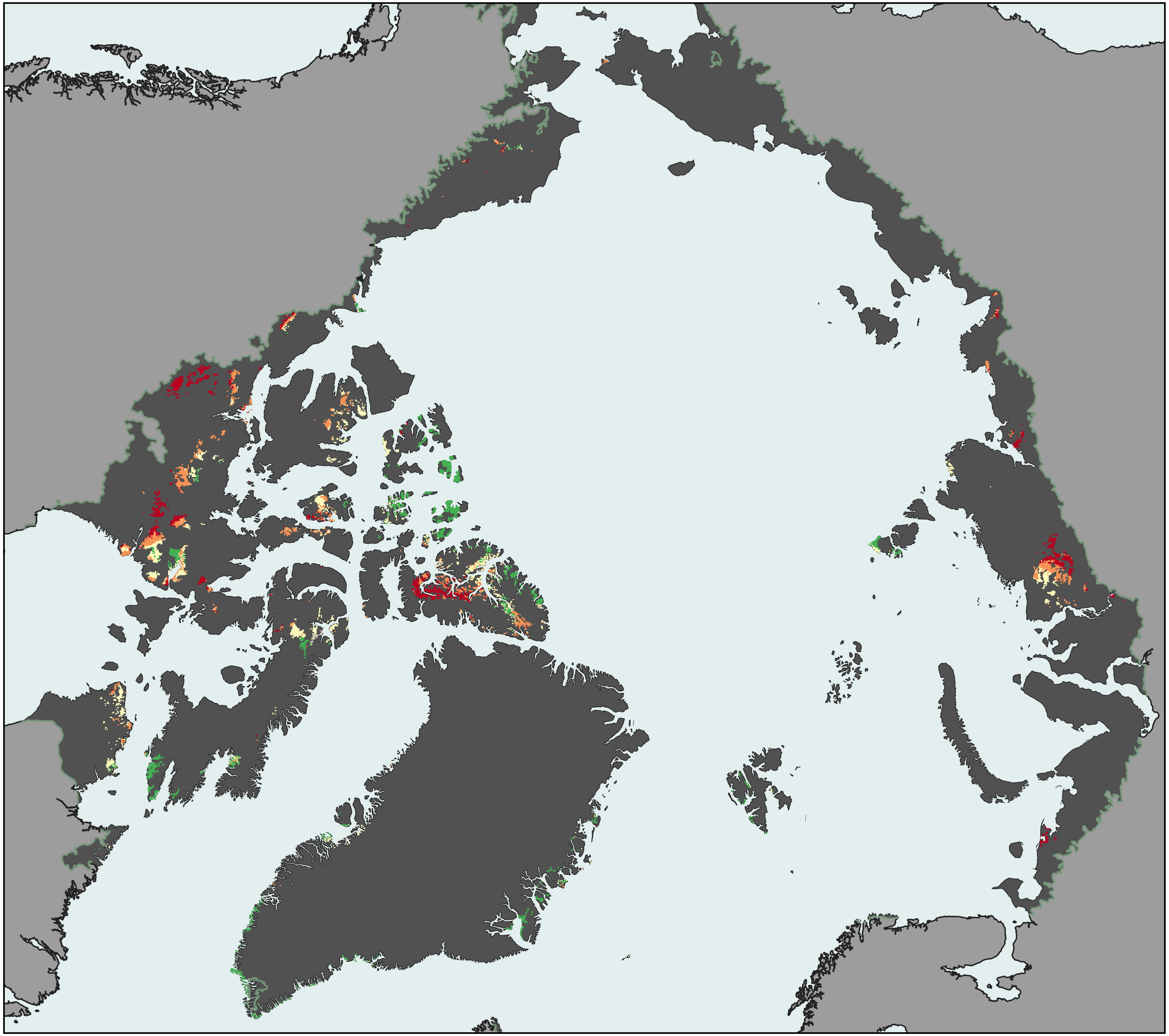

**Citation**

M. Reji Chacko, A.K.A. Goerens, J. Oehri, E. Plekhanova, and G. Schaepman-Strub. Will current protected areas harbour refugia for threatened Arctic vegetation types until 2050? Manuscript submitted.

**Project Director**

G. Schaepman-Strub

**Compilation & Cartography**

A.K.A. Goerens, M. Reji Chacko

University of Zurich[UZH]

URPP Global Change and Biodiversity
